# The population dynamic life histories of the birds and mammals of the world

**DOI:** 10.1101/2021.11.27.470200

**Authors:** Lars Witting

**Affiliations:** Greenland Institute of Natural Resources, Box 570, DK-3900 Nuuk, Greenland

**Keywords:** Life history, population dynamics, natural selection, bird, mammal, allometry

## Abstract

With life history traits determining the natural selection fitnesses of individuals and growth of populations, estimates of their variation are essential to advance evolutionary theory and ecological management during times of global change. As quantitative predictions improve with the completeness of models, and as data are usually incomplete or missing for most species, I use published data and inter-specific allometric extrapolations to estimate complete population dynamic life history models for birds and mammals with known body masses. This constructs models for 11,188 species of birds and 4,937 species of mammals, covering 27 life history and ecological traits per species. The estimates are used to illustrate natural selection mechanisms and explain a diverse range of population dynamic trajectories by the inclusion of population dynamic regulation. This provides a first step towards the construction of freely accessible and ready-to-use online population dynamic simulations covering all species of birds and mammals.

## 1 Introduction

With life history traits defining the fitness of individuals they provide a gateway to the secrets of natural selection, inspiring evolutionary studies for decades. And with the average fitness of individuals defining the growth of populations, life histories are equally essential for our ability to model and manage the dynamics of natural populations. Improving our skills in both areas are more critical now than ever, as the expanding demands of the growing human population challenge our living planet with climatic change and loss of habitats and biological diversity.

Solutions to the biodiversity and climate crises require political actions. As scientist we can help by integrating biological data into relevant measures, models, and analyses that are freely accessible and easy to interpret and use. This applies not only where data are available, but also by extrapolating our best scientific knowledge for use in critical cases where actions are needed but time and/or economy is too limited for data collection and detailed analysis.

We may use population dynamic data at face value, as in the Living Planet Index (LPI 2022) where timeseries of almost 40,000 populations measure the joint trend in the diversity of animal life on Earth. According to the LPI, vertebrates are on a global retreat with an average decline of about 70% from 1970 to 2018 across all monitored populations. Luckily, the LPI gives a false impression as the estimated decline follows from outliers, with 98.6% of the vertebrate populations across all systems showing no mean global trend (Leung et al. 2020). This highlights the need for species models to analyse the conservation implications of trends, identifying species and regions with need for action, as well as areas that improve.

Population trends have many causes, and their population biological implications are not always as straightforward as they seem. Successful trend analyses depend on realistic population models that can identify the underlying causes of timeseries. With this paper I initiates a project that uses population dynamic timeseries data to estimate first-approximation population dynamic models for all species of birds and mammals with an estimate of body mass. Apart from analysing the underlying causes of existing timeseries, a main goal is the construction of a global library of Bird and Mammal Populations (BMP) that is freely available for online simulations (at https://mrLife.org/bmp.htm). The BMP library allows for population dynamic analyses of all species, including the estimation of expected trajectories given habitats that fragment or improve.

The development of BMP includes three relatively independent stages, with this paper covering the first. Apart from externally imposed variation and trends, the population dynamics of a species is determined by its age-structured demography and population dynamic regulation by density dependent competition and natural selection. While the two regulating forces can be estimated from population dynamic timeseries, these data are generally insufficient for the estimation of the age-structured demography. Hence, in this first stage I use data from the literature to estimate age-structured life history models for all species with an estimate of mass. Having the age-structure demography I proceed in phase two with the estimation of population regulation for all species with timeseries data of a sufficiently high quality (following the approach in Witting 2021), before the estimates of regulation are extrapolated across species in phase three.

Instead of restricting my estimates to demographic parameters that are necessary for population dynamic simulations, I estimate a much broader set of life history traits where the feeding ecology and energetic physiology of each species are selected in a mutual balance with zero population dynamic growth. Being based on the population dynamic feedback selection of Malthusian relativity (Witting 1997, 2008, 2017a,b), these models have a strong theoretical basic, and a wide range of potential uses including analysis on evolutionary causality (Witting 2023).

### 1.1 Theoretical foundation

By analysing fitnesses in populations in relation to trade-offs and constraints, the success of classical life history theory (Charlesworth 1980; Roff 1992; Stearns 1992; Charnov 1993) is perhaps best described by it its ability to determine whether a species has evolved to a selection attractor, or not.

At first, this success seems to be somewhat of a paradox. This is because the constant relative fitnesses and frequency-independent selection that is applied in most traditional theory ignores the selection implications of the density and frequency dependent interactions that occur among individuals in natural populations. These interactions are typically competitive, and as they distribute the available resources differentially across individuals, they are directly influencing the relative fitnesses among variants (see e.g. Hardy and Briffa 2013). This makes the relative fitness of a variant dependent not only on the frequencies of the other life history variants in the population, but the frequency dependence is also density dependent as individuals encounter other individuals in competition more often at high population densities. How can a selection theory that does not include these basis rules of competition have any success with evolutionary predictions?

Population dynamic feedback selection takes the alternative game theoretical route (Maynard Smith and Price 1973; Maynard Smith 1982), using asymmetrical interactions (Parker 1974) and Continuously Stable Strategies (Eshel and Motro 1981; Taylor 1989; Christiansen 1991) to analyse life history evolution by the selection of the density-frequency-dependent competitive interactions among individuals in natural populations (including also a subset of trade-offs that cannot evolve). Where most game theory deal with differences in behavioural strategies of competition, population dynamic feedback selection focusses mainly on the interactive asymmetry obtained by life history traits like body mass, group size, the male/female division of labour, and the associated sexual reproduction attraction of two genders. These interactive qualities are inversely related to population dynamic growth by the quality-quantity trade-off (Smith and Fretwell 1974; Stearns 1992), the costs of resource sharing, the two-fold cost of males (Maynard Smith 1971), and the two-fold cost of meiosis (Williams 1975). Their interactive selection is thus controlling the growth, abundance, and level of interference of the population, creating a population dynamic feedback that selects for energetically balanced life histories (Witting 1997, 2002, 2008).

Where classical life history theory often is verified by its ability to mimic fitness distributions in natural populations, feedback selection is tested mainly by its ability to predict evolution in time, including the observed differentiation in life histories among species and higher taxa. This involves *i*) the prediction of lifeforms from molecular replicators over prokaryote and protozoa like self-replicating cells to multicellular animals (Witting 2017b), *ii*) the associated prediction of replicating units from asexual replicators over sexually reproducing pairs and cooperative breeders to fully evolved eusocial colonies (Witting 1997, 2002, 2007), and *iii*) the deduction of the inter-specific body mass allometries (Witting 1995, 2017a) that I use in the present paper to extrapolate life history models across birds and mammals.

The difference between the frequency-independent and density-frequency-dependent fitnesses seems to place classical life history theory and population dynamic feedback selection in direct opposition to one another. Yet, it is actually possible to consolidate the two frameworks into a coherent whole by treating the classical method as an instant description of current selection, and population dynamic feedback selection as a description of natural selection through time, including predictions of life history variation between species. This is because the density-frequency-dependence is frozen at instant moments in time, where the population densities and levels of interference are given, and the distribution of variants and relative fitnesses are constant.

The distributions of fitnesses in natural populations at given points in time may thus be analysed relatively easily by the mathematical framework of constant relative fitnesses to estimate the current outcome of natural selection in relation to the current physiological and ecological trade-offs experience by the species. Yet—as the population density, level of interference, composition of variants, and relative fitnesses start to change with evolution—the frequency-independent framework is usually too rigid for realistic predictions of evolutionary variation across species and time. When attempted, distorted predictions are often obtained (Witting 1997, 2008).

A distortion is evident in allometric theory, where the frequency-independent framework of the metabolic theory of ecology (West et al. 1997, 1999; Brown et al. 2004; Brown and Sibly 2006) predicts an increase in the population dynamic growth rate (*r*) and carrying capacity (*k*) with a selection increase in mass (Witting 2018, 2023). These traits, however, are typically not increasing but declining with an increase in mass across natural species (Fenchel 1974; Damuth 1981, 1987), questioning the evolutionary relevance of the metabolic theory of ecology.

By explicitly analysing the selection of density dependent interactive competition, population dynamic feedback selection provides the first and so far only prediction where the inter-specific allometric exponents of the physiological and ecological traits in Table 1 are explained by the natural selection of metabolism and mass (Witting 1995, 2017a), including the observed decline in *r* and *k* with a selection increase in mass. This prediction includes not only well-known Kleiber (1932) scaling with typical ±1*/*4 and ±3*/*4 exponents in terrestrial taxa that forage in predominantly two spatial dimensions, but also corresponding ±1*/*6 and ±5*/*6 exponents in pelagic species that forage in three spatial dimensions (Witting 1995), and *i*) allometric transitions from prokaryotes to protozoa and multicellular animals (Witting 2017a,b), *ii*) a curvature in the metabolic allometry of placentals (Witting 2018), and *iii*) numerical scaling differences in body mass trajectories of the fossil record predicting biological evolution over millions and billions of years (Witting 2020).

**Table 1:** Top table: The number of birds (Ave) and mammals (Mam) with data estimates across traits. Bottom table: The body mass allometry exponents predicted by population dynamic feedback selection; see mathematical deductions in Witting (1995, 2017a) or graphical deduction in Witting (2023) [*t*_*x*_ ∈ {*t*_*p*_, *t*_*j*_, *t*_*s*_, *t*_*m*_, *t*_*r*_, *t*_*g*_, *t*_*l*_}]. These exponents depend on the ecological packing of home ranges in one, two, or three spatial dimensions (*d*), as listed in Table 2.

| | $w$ | $w_0$ | $w_j$ | $w_f$ | $\beta$ | $\underline{\beta}$ | $t_p$ | $t_j$ | $t_s$ | $t_m$ | $t_l$ | $m_b$ | $m_f$ | $m_i$ | $p_{ad}$ | $N$ | $h$ |
| --- | --- | --- | --- | --- | --- | --- | --- | --- | --- | --- | --- | --- | --- | --- | --- | --- | --- |
| Ave. | 11188 | 774 | 207 | 155 | 76 | 470 | 2237 | 1819 | 1210 | 681 | 1667 | 6979 | 1765 | 0 | 908 | 1438 | 249 |
| Mam. | 4937 | 1868 | 1059 | 101 | 57 | 713 | 2217 | 2040 | 1999 | 935 | 2614 | 3511 | 2146 | 651 | 310 | 1111 | 456 |

Table 1: Top table: The number of birds (Ave) and mammals (Mam) with data estimates across traits.
| | $w$ | $\beta$ | $t_x$ | $m$ | $R$ | $l_m$ | $l_r$ | $\epsilon$ | $\alpha$ | $N$ | $b$ | $\epsilon_n$ | $h$ | $h_o$ | $I$ |
| --- | --- | --- | --- | --- | --- | --- | --- | --- | --- | --- | --- | --- | --- | --- | --- |
| Exp. | 1 | $-\frac{1}{2d}$ | $\frac{1}{2d}$ | $-\frac{1}{2d}$ | 0 | 0 | 0 | $\frac{2d-1}{2d}$ | 1 | $\frac{1-2d}{2d}$ | $\frac{1}{2d}$ | 0 | 1 | $\frac{1}{2d}$ | 0 |

This selection of metabolism, body mass, and allometries follows from a population dynamic feedback where the interactive foraging in overlapping home ranges generates net energy for reproduction, and thus drives the population dynamic growth that determines the population dynamic equilibrium abundance, which in term defines the home range overlap and density dependent level of interference competition that selects net energy into mass, as the larger-than-average individuals monopolise resources during interactive encounters. The selection is defined and constrained by the physiological energetics of individuals, the population dynamic growth, and the resulting foraging ecology where individual opportunities are constrained by the spatial dimensionality of the habitat, the abundance of the population, and the trait-values of the other individuals in the population. This involves a mass-rescaling selection where Kleiber scaling is selected as a secondary response that maintains the net energy of the organism during the selection of mass (Witting 2017a).

It is the life history and ecological traits that are linked to one another in the population dynamic feedback cycle that I estimate in this study. The estimated models are thus directly compatible with the energy flow in population dynamic feedback selection, except that the estimates focus only on the average trait-values, and not on the intra-population differentiation that defines natural selection. For insights on the latter you may consult theoretical studies like Witting (1997, 2002, 2008, 2017a,b).

A main purpose of my paper is to construct the diversity of models that cover the population dynamic life history variation across all species of birds and mammals. Yet, I will also use the intra- and inter-taxa variation of birds, placentals, marsupials, and bats to illustrate that the physiology, demography, and ecology of natural species are selected in a mutual balance that follows the expectations of population dynamic feedback selection. And at the end of the paper, I illustrate how the single species models are easy to extend into simulations that explain a diverse set of population dynamic timeseries by the inclusion of population dynamic regulation.

## 2 Methods

To estimate the missing life history traits of the different species I combine large datasets published over the past couple of decades with inter-specific extrapolations by allometric correlations reflecting population dynamic feedback selection. Following this method, the minimum requirement for a species model is an estimate of body mass.

To extrapolate life history models across species, I combine the diverse data into a single dataset, and use standard calculations to transform some of the data into common parameters. The data are then checked for outliers and combined with allometric calculations to estimate the missing values for all species with a data-based estimate of body mass.

All of my life history estimates follow the model structure that Witting (2017a,b) used to deduce the exponents of inter-specific allometries from the natural selection of metabolism, body mass, and the life history and population dynamic foraging ecology as a whole. For a detailed description of the underlying population dynamic feedback selection, you will have to consult these papers, as I restrict myself to the estimation of the underlying population level parameters. Some basic demographic relations are described in Appendix A, with the overall model divided into the four components of individual growth, demography, life history energetics, and population ecology. The four sub-sets of parameters are introduced below.

### 2.1 Individual growth

The growth of an individual with age (*a*) is described by

*t*_*p*_: the gestation period in mammals, and incubation period in birds;

*t*_0_: the age at birth in mammals, and age at hatching in birds, *t*_0_ = 0;

*t*_*f*_: the age at maximum growth, i.e., infliction point;

*t*_*j*_: the age at weaning in mammals, and age at fledging in birds;

*t*_*d*_: the age at independence from parents, *t*_*j*_ ≤ *t*_*d*_ *< t*_*m*_ with *t*_*m*_ being the age at reproductive maturity, assuming complete parental care for *a < t*_*d*_, and no care for *a* ≥ *t*_*d*_;

*w*: the average body mass of an adult individual;

*w*_*x*_: the mass at *t*_*x*_ = *t*_0_, *t*_*f*_, *t*_*j*_ or *t*_*d*_, with relative mass 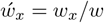.

Embryotic growth from age *t*_*p*_ to *t*_0_ is calculated by the Gompertz (1832) growth curve, following the regularities estimated by Ricklefs (2010), where the asymptote 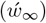 of the Gompertz growth function 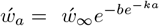 is 2.5 times 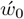 for embryotic growth, and the growth rate 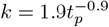 is approximately inversely proportional to the gestation/incubation period. This implies the following embryotic growth

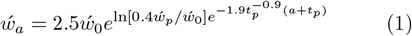

with age −*t*_*p*_ ≤*a <* 0 given in days and relative mass at −*t*_*p*_ being 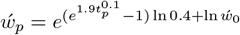.

Growth after birth/hatching is estimated by the Richards (1959) growth function, with species specific data-estimates of the inflection point obtained by least-square fitting to growth data obtained from the literature [supplementary information (SI) on inflection]. The growth function

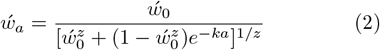

passes through 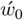 at *a* = *t*_0_ = 0 and 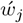 at *a* = *t*_*j*_, with 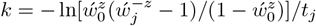 and a relative adult mass of unity 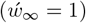. The infliction point 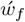 relative to adult mass is calculated given the fitted or extrapolated estimates of the shape parameter *z*, with 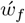 increasing from zero to 0.5 when *z* increases from −1 to 1.

There is only little information available on *t*_*d*_ and 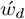 (but see e.g. Millar et al. 1986; Lloyd 1987; Johansson et al. 2021). While I treat independence as a knife-edge parameter, real independence is a gradual development. Independence is often quite late compared to weaning in mammals, and often quite close to fledging in birds [with median 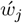 being 0.33 (cv:0.77) and 0.84 (cv:0.24) in mammals and birds for the data in the present study]. With no clear data estimates, I assume that parents invest at least 80% in the mass of offspring (either directly, or indirectly by protecting the foraging area of offspring). The relative mass at independence is thus 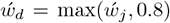, with *t*_*d*_ being the solution to eqn 2 at 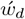.

### 2.2 Demography

The demographic traits include

*p*_*ad*_: the annual survival probability of adults (geometric mean across adult age-classes);

*q*_*ad*_: the annual mortality of adults, *q*_*ad*_ = 1 − *p*_*ad*_; *t*_*s*_: female age at sexual maturity;

*t*_*m*_: female age at reproductive maturity, *t*_*m*_ = *t*_*s*_ + *t*_*p*_;

*t*_*r*_: the expected reproductive period for females that survive to *t*_*m*_, estimated as *t*_*r*_ = 1*/q*_*ad*_ by eqn 12;

*t*_*g*_: generation time as the average age of reproduction, as *t*_*g*_ = *t*_*m*_ + *p*_*ad*_*/q*_*ad*_ by eqn 14;

*t*_*l*_: the maximum lifespan of an individual;

*l*_*m*_: the probability that a new-born survives to *t*_*m*_, as

*l*_*m*_ = 2*/R* by eqn 3;

*l*_*r*_: the probability that an adult survives the expected reproductive period, 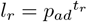;

*m*: the average number of offspring produced per female per year, i.e., *m* = *m*_*b*_ *m*_*f*_ where *m*_*b*_ is the average brood size and *m*_*f*_ the average brood frequency, with *m*_*f*_ being the inverse *m*_*f*_ = 1*/m*_*i*_ of the average brood interval *m*_*i*_; and

*R*: expected lifetime reproduction as the average number of offspring produced over the expected reproductive period of females that survive to *t*_*m*_, estimated as *R* = *t*_*r*_*m*.

These parameters are estimated for species that are assumed to be naturally selected around a population dynamic equilibrium with a per-generation growth rate of unity

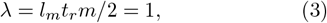

assuming an even sex ratio.

### 2.3 Life history energetics

Traits that link the demographic parameters to the energetics of the organism include

*w*_*E*_: body mass as combustion energy (SI unit J); obtained as *w*_*E*_ = *c*_*w*→*d*_*c*_*d*→*E*_*w*, with *w* being mass in grams, *c*_*w*→*d*_ the conversion of wet organic matter to dry [0.40 for birds (Mahoney and Jehl 1984), and 0.35 for mammals (Prothero 2015)], and *c*_*d*→*E*_ the conversion of dry matter to energy [22.6 kJ/g from Odum et al. (1965) and Griffiths (1977)];

*<u>β</u>*: basal mass-specific metabolism (SI unit W/g);

*β*: field mass-specific metabolism (SI unit W/g) also the pace (SI unit Hz) of the metabolic work carried out per unit body mass;

*α*: the handling of net resource assimilation (resource handling in short, SI unit J), defined as the net energy that is generated per metabolic pace (estimated as *α* = *E/β*). Handling is a joint parameter 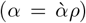 of intrinsic handling 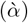 multiplied by the density of the resource (*ρ*);

*E*: the net assimilated energy of adults is the product (*E* = *αβ*; SI unit W) of resource handling (*α*) and metabolic pace (*β*). Following the selection attractor of sexually reproducing pairs (Witting 2002), half of the join net energy (2*E*) of the female/male unit is allocated to reproduction, while the other half is used in intra-specific interactions. Net energy is estimated as 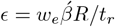 from eqn 4; and *E*_*g*_: average gross energy *E*_*g*_ = *wβ* + *E/*2 (SI unit W; given as a relative measure 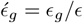) is total field metabolism plus net energy allocated to reproduction. The latter is *E/*2 as the reproducing unit uses half of their net energy in reproduction, while the other half that is used in intra-specific interactions is included here in average field metabolism.

Net energy defines lifetime reproduction by the number of fully growth offspring that it produces. Assuming complete parental investment until *t*_*d*_, we have

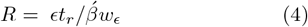

where

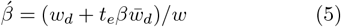

is a scaling parameter that scales *w*_*E*_ to the mass at independence (*w*_*d*_) and accounts for the energy that is metabolised by the offspring during parental care (*t*_*e*_ = *t*_*p*_ + *t*_*d*_), with 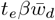 being the energy that is metabolised by the offspring during *t*_*e*_, assuming constant mass-specific metabolism (*β*) and an average size

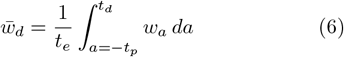

that is calculated by the growth model of eqns 1 and 2, with −*t*_*p*_ being the negative age at fertilisation.

### 2.4 Population ecology

Important population ecological components include

*N*: population density (abundance), as the number of animals per km^2^;

*b*: the standing biomass of the population, as *b* = *wN* in kg/km^2^;

*E*_*n*_: the energy consumed by the population, calculated as *E*_*n*_ = *E*_*g*_*N* (in W/km^2^);

*h*: the average home range of an individual (in km^2^);

*h*_*o*_: the average home ranges overlap *h*_*o*_ = *hn*, given as home range divided by the available of space (1*/N*) per individual;

*v*: the frequency of competitive encounters per individual per unit home range overlap, given as a relative measure *v* ∝ *v*_*f*_ */l*_*f*_, where *l*_*f*_ ∝*h*^1*/d*^ is the length of foraging tracks (proportional to the *d*th root of the home range in *d*-dimensions) and *v*_*f*_ ∝*β*_*β*_*w*^1*/*2*d*^ the average foraging speed obtained from allometric correlations where *β*_*β*_ is the intercept of the metabolic allometry *β* = *β*_*β*_*w*^−1*/*2*d*^ (Witting 1995, 2017a); and

*I*: intra-specific interference competition per individual, calculated as a relative measure *I* ∝ *h*_*o*_*v* ∝ *Nhv* from the overlap in home ranges and the frequency of encountering other individuals per unit home range overlap (Witting 1995, 2017a). These *I* estimates are rescaled for a log value (*℩* = ln *I*) of unity at the median across all species of birds.

### 2.5 Data

My study extends on the data used by Witting (2023), with all literature estimates treated as raw data. My broad definition of data includes the traits of a species that are calculated from the raw data of that species.

I use the BirdLife (2022) taxonomy for birds, and MDD (2023) taxonomy for mammals, with some subspecies with separate body mass estimates categorised as species. Body mass data were obtained from several sources including Smith et al. (2004), Dunning (2007), Jones et al. (2009), Weisbecker et al. (2013), Myhrvold et al. (2015), Tobias et al. (2021), and Herberstein et al. (2022). Demographic parameters on reproduction, time periods, and growth were achieved from many datasets including Jetz et al. (2008), De Magalhães and Costa (2009), Jones et al. (2009), and Myhrvold et al. (2015), with growth curve data and survival rates obtained from an independent literature search, including survival estimates from McCarthy et al. (2008), DeSante and Kaschube (2009), Ricklefs et al. (2011), del Hoyo et al. (1992–2011), and Wilson and Mittermeier (2009– 2014). Ecological parameters included population densities from Damuth (1987) and Santini et al. (2018), and home range areas from Tucker et al. (2014), Tamburello et al. (2015), and Nasrinpour et al. (2017) with an independent literature search for marine mammals and bats. Basal metabolic rates were obtained primarily from McNab (2008) for mammals and from McKechnie and Wolf (2004) for birds, with field metabolism for both taxa provided by Hudson et al. (2013).

Altogether, 110,105 trait values were obtained from 614 literature sources. Some of these data are the same, and I put more weight on commonly agreed estimates by using the average trait values of the available raw data for each trait per species. This resulted in 58,548 species specific raw data, with Table 1 listing the number of species with raw data for different traits.

These data are distributed relatively evenly across the taxonomic groups considered separately in this paper. For the traits in Table 1, there were on average raw data for 17% of the bird species and 23% of the mammalian species, with the latter including 25% for placentals, 31% for marsupials, and 15% for bats. These data, except for body mass estimates, were checked for outliers (see outlier SI), with a total of 474 outliers removed.

I list the literature references for all raw data (follow the letter codes in the SI on estimates and SI on data references), and for the underlying raw data when derived traits are calculated from one or two underlying traits with raw data estimates. Higher-level data that are calculated from more traits are listed as data but with no specific reference. Traits that could not be calculated from the raw data of a species had missing values that were estimated by inter-specific extrapolations.

### 2.6 Estimating missing values

I estimated missing values by inter-specific body mass allometries and invariances following the allometric model of Witting (2017a), using the estimation sequence in Appendix B.

#### Estimation level

As my data come from many different sources, and as they have not been collected as a random sample across the different species, they are phylogenetically unevenly distributed. This means that the uncertainties of the missing value estimates will increase with the phylogenetic distance to the nearest species with data. To capture this uneven distribution of uncertainties, I use a missing parameter estimator that differentiates between the different phylogenetic levels. This provides separate estimates of uncertainty for the different parameters at the different phylogenetic estimator levels. The uncertainty distribution is visualised for all parameter estimates across all species by a colour code, where data are presented in black and missing parameter estimates at the genus, family, order, and class levels are presented in blue, green, yellow, and red.

Missing values were estimated at the different taxonomic levels dependent upon the available data. The precision of an allometric estimate will generally increase with the number of data and decline with taxonomic level from genus over family and order to class. With the number of data points for any trait increasing with taxonomic level, we expect a trade-off where an estimator based on few data at a low taxonomic level will, at some point, provide a better estimate than an estimator with many data at a higher taxonomic level. To determine the data limits—where lower-level estimators are preferred over higher-level estimators with more data—I constructed a hierarchical estimator that used the lowest taxonomic level with a given number (*n*_*d*_) of data points. This number was estimated by a numerical minimization of the sum of squares of the difference between raw data and their predicted values, with the predicted data values of a species being predicted from the raw data of all other species.

Given the optimal missing value estimation of eqns 7 to 9 below, the minimization of the sum of squares between the raw data and their predicted values determined that the more precise estimates of missing values were obtained with a *n*_*d*_ parameter around unity. Taxonomic proximity was thus prioritized over sample size, with missing values being calculated at the lowest taxonomic level with one or more raw data of the required parameters. Genus (*g*) was prioritized over family (*f*), family over order(*o*), and order over class (*c*).

#### Estimator optimization

The missing values of mass-dependent parameters were calculated as allometric functions of mass. The age of reproductive maturity, e.g., was calculated as

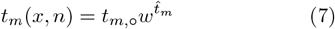

with *x* denoting the lowest taxonomic level with raw data, and *n* the number of data points at that level. The allometric exponent 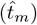 and intercept (*t*_*m*,°_) were estimated as a joint function

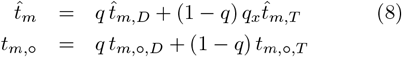

of the theoretical expectations from Witting (1995, 2017a) [subscript *T*; e.g. 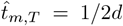 from Table 1 with ecological dimensionality (*d*) from Table 2] and the *n* data points at the *x*th taxonomic level [subscript *D*, with 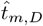 and *t*_*m*,°,*D*_ being point estimates from a linear regression on double logarithmic scale]. The weight

**Table 2:** The spatial dimensionality (d) of the ecological packing of home ranges is set to depend on taxonomic level, following the classification in Witting (2017a).

| Class | Order | Family | $d$ |
| --- | --- | --- | --- |
| Aves | - | - | 2 |
| Mammalia | - | - | 2 |
| Mammalia | Primates | - | 3 |
| Mammalia | Carnivora | - | 2 |
| Mammalia | Carnivora | Otariidae | 3 |
| Mammalia | Carnivora | Odobenidae | 3 |
| Mammalia | Carnivora | Phocidae | 3 |
| Mammalia | Cetacea | - | 3 |

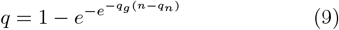

of the data estimate increases monotonically from zero to one as a function of *n*, except for *n* ≤ 3 cases where the exponent 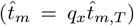 was proportional to the theoretical expectation and the *t*_*m*,°_ intercept of eqn 7 was the average allometric intercept for the *n* species given the 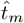 exponent. Having no theoretical expectation for 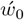 and 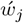, their theoretical values were replaced with exponents estimated by linear regression across all raw data for birds, placentals (minus bats), marsupials, and bats (estimates varying from −0.07 to 0.22).

The three parameters *q*_*g*_, *q*_*n*_, and *q*_*x*_ were estimated separately for all mass dependent parameters. This was done by a numerical minimization of the sum of squares (across species with data) of the difference between their raw data and the predicted values by the estimators at the lowest taxonomic levels, given the data of all the other species.

Invariance estimators were used only for lifetime reproduction (*R*) and the inflection parameter *z*. These were estimated as the average of the data across the lowest taxonomic level with data for that parameter.

Given estimators at the lowest taxonomic level with data, the optimal values of the parameters for eqns 8 and 9 are listed in Table 3. They follow a general pattern where missing parameters are calculated from the theoretical exponents for estimators with few available data. With an increased number of data there often is a steep transition to a fully empirically based estimator, with the average values of the empirically estimated exponents listed in Table 3, for estimators at the genus (blue), family (green), order (yellow), and class (red) level.

**Table 3:** Missing value estimation. The *q*_*g*_, *q*_*n*_ and *q*_*x*_ parameters of eqns 8 and 9 that give the most precise estimation of missing values for the different mass-dependent traits. 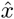 is the average, and *σ* the standard deviation, of the data estimated allometric exponents at the different taxonomic levels (blue:genus; green:family; yellow:order; red:class). The precision of an estimated missing value is given by a coefficient of variation (cv) that is estimated from the missing value predictions of n data points.

| | $w_0$ | $w_j$ | $\beta$ | $\underline{\beta}$ | $t_m$ | $t_l$ | $t_p$ | $t_j$ | $m$ | $p_{ad}$ | $q_{ad}$ | $N$ | $h$ |
| --- | --- | --- | --- | --- | --- | --- | --- | --- | --- | --- | --- | --- | --- |
| $q_g$ | 3.53 | 4.61 | 0.16 | 3.99 | 3.39 | 0.05 | 2.60 | 0.08 | 3.39 | 0.06 | 0.26 | 0.81 | 0.26 |
| $q_n$ | 8.1 | 10.7 | 30.6 | 27.0 | 17.5 | 100.0 | 19.0 | 46.5 | 29.9 | 54.6 | 19.0 | 83.9 | 28.0 |
| $q_x$ | 0.33 | 0.33 | 1.05 | 1.02 | 0.47 | 0.78 | 0.33 | 0.51 | 0.33 | 0.42 | 0.85 | 0.62 | 0.99 |
| $\hat{x}$ | -0.34 | -0.15 | -0.16 | -0.11 | 0.02 | 0.16 | 0.03 | 0.14 | 0.06 | -0.11 | 0.09 | -0.34 | 0.95 |
| $\sigma$ | 0.47 | 0.71 | 0.79 | 0.65 | 0.98 | 1.18 | 0.39 | 0.58 | 1.05 | 1.05 | 1.22 | 3.98 | 3.45 |
| $\hat{x}$ | -0.30 | -0.12 | -0.26 | -0.27 | 0.13 | 0.13 | 0.07 | 0.15 | -0.08 | 0.08 | -0.14 | -0.39 | 0.86 |
| $\sigma$ | 0.32 | 0.54 | 0.29 | 0.27 | 0.39 | 0.77 | 0.20 | 0.28 | 0.60 | 0.25 | 0.44 | 1.44 | 1.98 |
| $\hat{x}$ | -0.20 | -0.13 | -0.29 | -0.30 | 0.24 | 0.21 | 0.13 | 0.20 | -0.16 | 0.10 | -0.23 | -0.37 | 0.92 |
| $\sigma$ | 0.30 | 0.43 | 0.20 | 0.22 | 0.37 | 0.47 | 0.10 | 0.17 | 0.80 | 0.07 | 0.24 | 0.56 | 0.89 |
| $\hat{x}$ | -0.16 | -0.09 | -0.33 | -0.31 | 0.22 | 0.22 | 0.18 | 0.24 | -0.16 | 0.08 | -0.21 | -0.55 | 1.06 |
| $\sigma$ | 0.06 | 0.05 | 0.03 | 0.06 | 0.01 | 0 | 0 | 0.03 | 0.02 | 0 | 0.01 | 0.16 | 0.03 |
| cv | 0.14 | 0.15 | 0.14 | 0.12 | 0.14 | 0.24 | 0.05 | 0.09 | 0.13 | 0.12 | 0.21 | 0.87 | 1.04 |
| n | 2117 | 985 | 682 | 644 | 2446 | 3526 | 3653 | 3133 | 3112 | 695 | 695 | 1742 | 337 |
| cv | 0.24 | 0.21 | 0.15 | 0.15 | 0.20 | 0.25 | 0.11 | 0.18 | 0.23 | 0.17 | 0.25 | 0.95 | 1.12 |
| n | 459 | 227 | 477 | 462 | 507 | 629 | 728 | 647 | 624 | 376 | 376 | 712 | 289 |
| cv | 0.38 | 0.25 | 0.23 | 0.20 | 0.24 | 0.31 | 0.20 | 0.31 | 0.30 | 0.20 | 0.35 | 1.38 | 1.35 |
| n | 50 | 45 | 66 | 67 | 46 | 45 | 61 | 57 | 58 | 69 | 69 | 49 | 42 |
| cv | 2.26 | 0.26 | 0.33 | 0.26 | 0.48 | 0.33 | 0.12 | 0.52 | 0.25 | 0.10 | 0.32 | 1.04 | 2.70 |
| n | 6 | 7 | 9 | 9 | 5 | 5 | 4 | 2 | 2 | 7 | 7 | 12 | 8 |

The average precision (cv) of the missing parameter estimates are also listed in Table 3. Excluding the estimator for abundance (*N*) and home range (*h*), the precisions are generally acceptable for all estimators. The average cv is 0.23, with a range that varies from 0.05 to 2.70. Precision is generally declining with the taxonomic level of the estimator, with the average cv being 0.20 at the genus level, 0.32 at the family level, 0.41 at the order level, and 0.77 at the class level.

A cross validation, which shows how well the different raw data points are predicted by the inter-specific extrapolations that exclude the estimated data points, is shown in Fig. 1. There is a close to linear dependence for all traits, yet some traits tend to be overestimated for small masses (see e.g. residual plots for juvenile mass and abundance).

**Figure 1:**
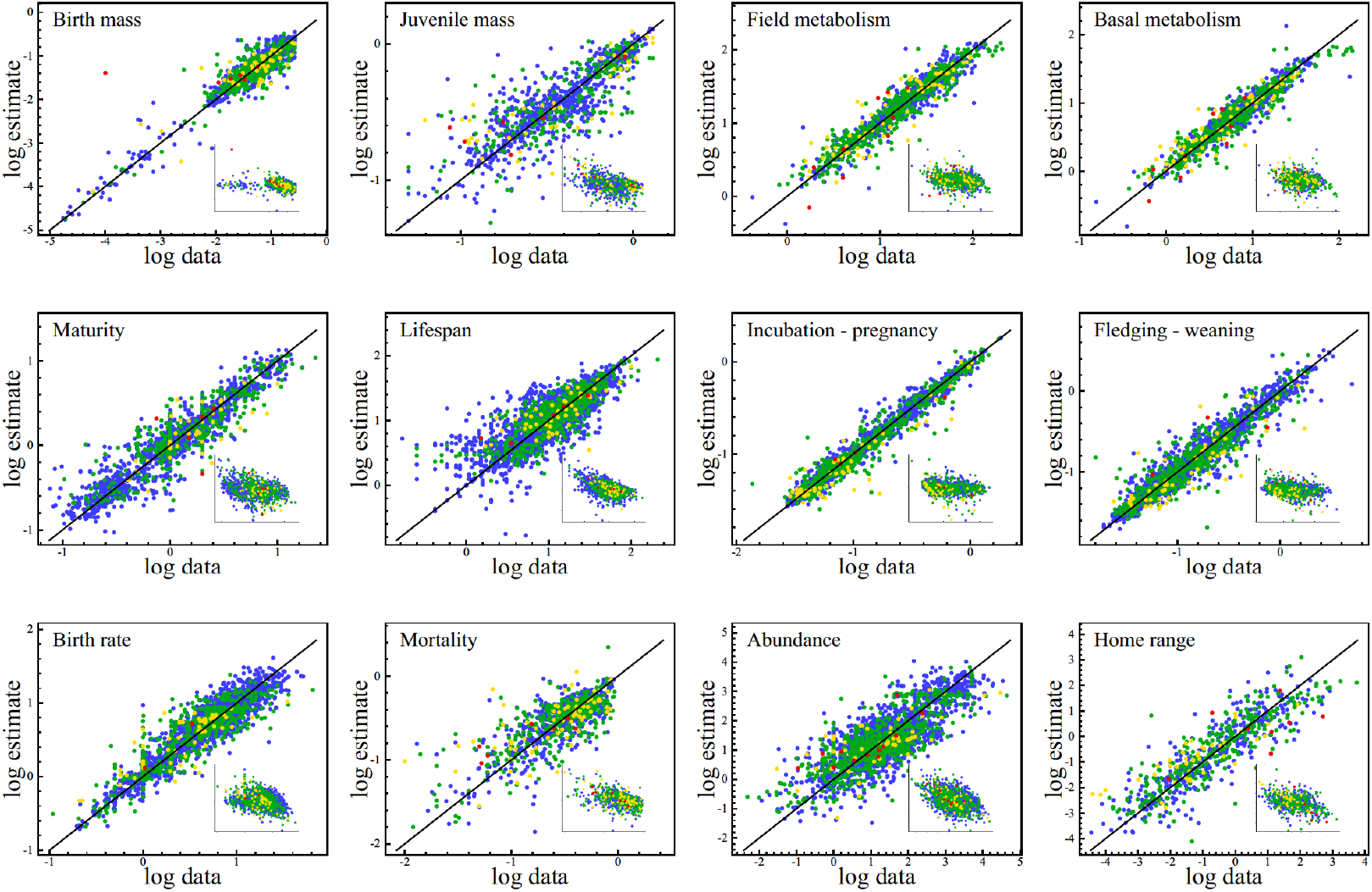
Data estimation. The estimation of missing parameters by allometric correlations was cross-validated and optimised by the ability to predict the actual data; as illustrated here by the relationships between the data and their estimates on double logarithmic scale. Residuals are shown in insert plots, and the precision (cv) of the estimates are given in Table 3. Estimator levels: genus (blue), family (green), order (yellow), and class (red); with points of the latter sitting on top of the former.

## 3 Results

### 3.1 Estimation level & space

Fig. 2 shows the proportions of estimation levels for the different parameter. Mammals have a larger fraction of data than birds for most traits, yet even for birds 85%, 93%, 92%, 88%, 74%, and 86% of the estimates for *m, N* (and *b*), *t*_*p*_, *t*_*j*_, *t*_*m*_ and *t*_*l*_ are at or below the family level. The corresponding values are 99%, 86%, 99%, 99%, 98%, and 96% for mammals, where 42%, 22%, 45%, 41%, 36%, and 45% of the estimates are data. For mass-specific metabolism 5% and 15% of the estimates for birds and mammals are data, and 84% and 95% of the estimates are at or below the family level. For *t*_*r*_ = 1*/q*_*ad*_, 8% and 4% of the estimates for birds and mammals are data, and 88% and 77% are at or below the family level. The mass at birth 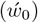 and weaning 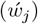 have a strong data base in mammals (37% & 21% are data), but less so in birds especially for mass at fledging (7% & 2% are data).

**Figure 2:**
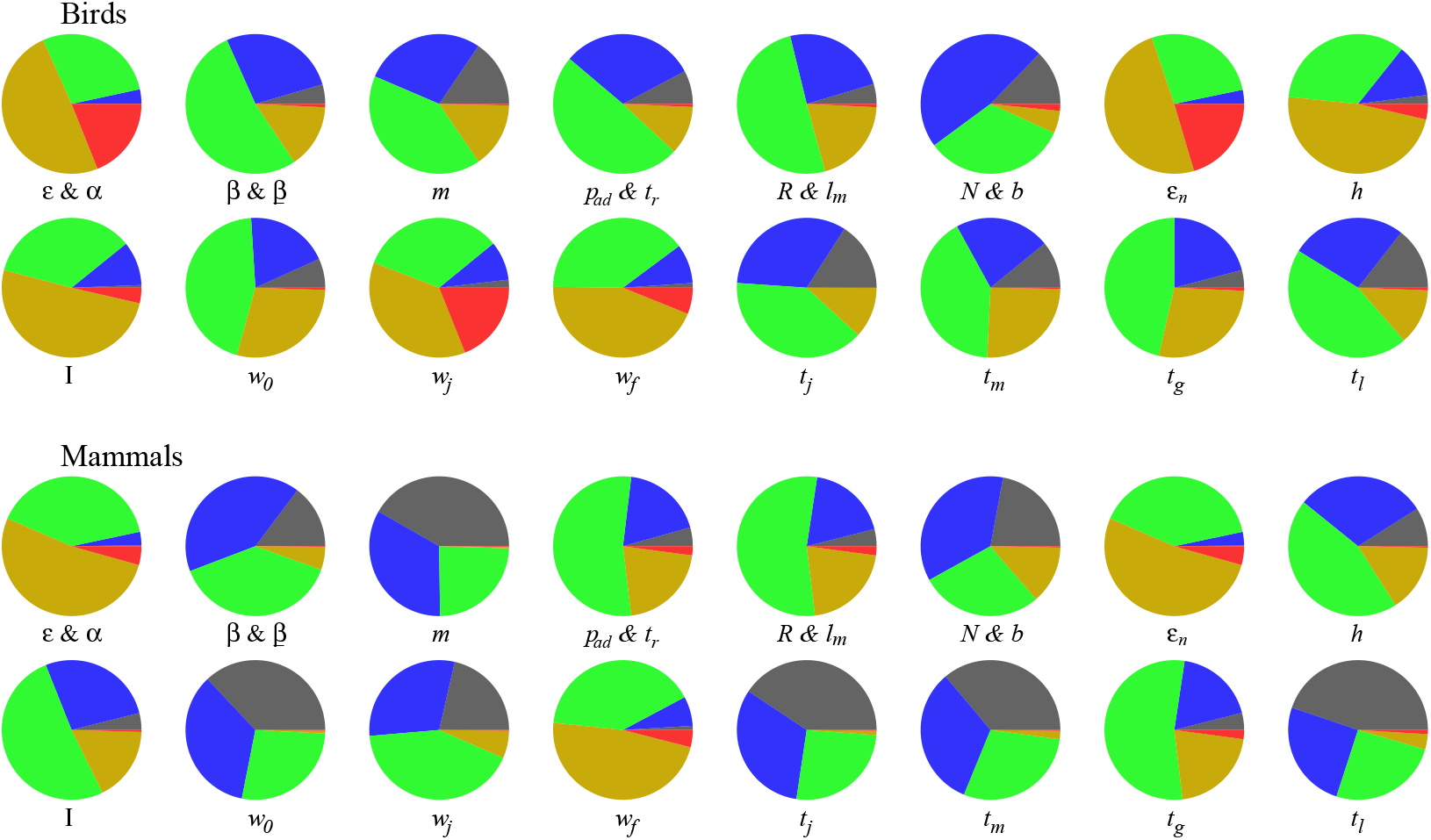
Estimator distributions for 11,188 species of birds, and 4,937 species of mammals. Estimator levels: data (black), genus (blue), family (green), order (yellow), and class (red).

Traits that are less supported by data are typically calculated from at least two independent parameters, with estimator levels given by the parameter that is estimated at the highest taxonomic level. These traits include *E, α, R, l*_*m*_, *E*_*n*_, *I*, and *t*_*g*_. Yet, even net energy (*E*) and resource handling (*α*)—which are calculated from seven underlying parameters—have 32% and 44% of the estimates in birds and mammals at or below the family level.

The estimated life history variation is plotted as allometric functions of body mass for birds and placentals in Fig. 3 and marsupials and bats in Fig. 4. The noncrossing black lines in the plots connect data, forming trees that illustrate the parameter space of the data. The estimated parameters for individual species are plotted by dots, with the taxonomic level of the estimator indicated by colour. The dots and lines show that the extrapolation of parameters across species are contained within the overall parameter space of the data. The general overlap between estimates and data holds especially for traits with many data, where the black trees cover almost the complete parameter space of the estimates. Yet, narrow extrapolation areas exist, like the line of order-level dots (yellow) for the allometric scaling of metabolism (*β*) in larger placentals. The parameters with fewer data have large data free areas that are dominated by especially order and family level estimators. But even these extrapolation areas tend to be contained within the space of scattered data.

**Figure 3:**
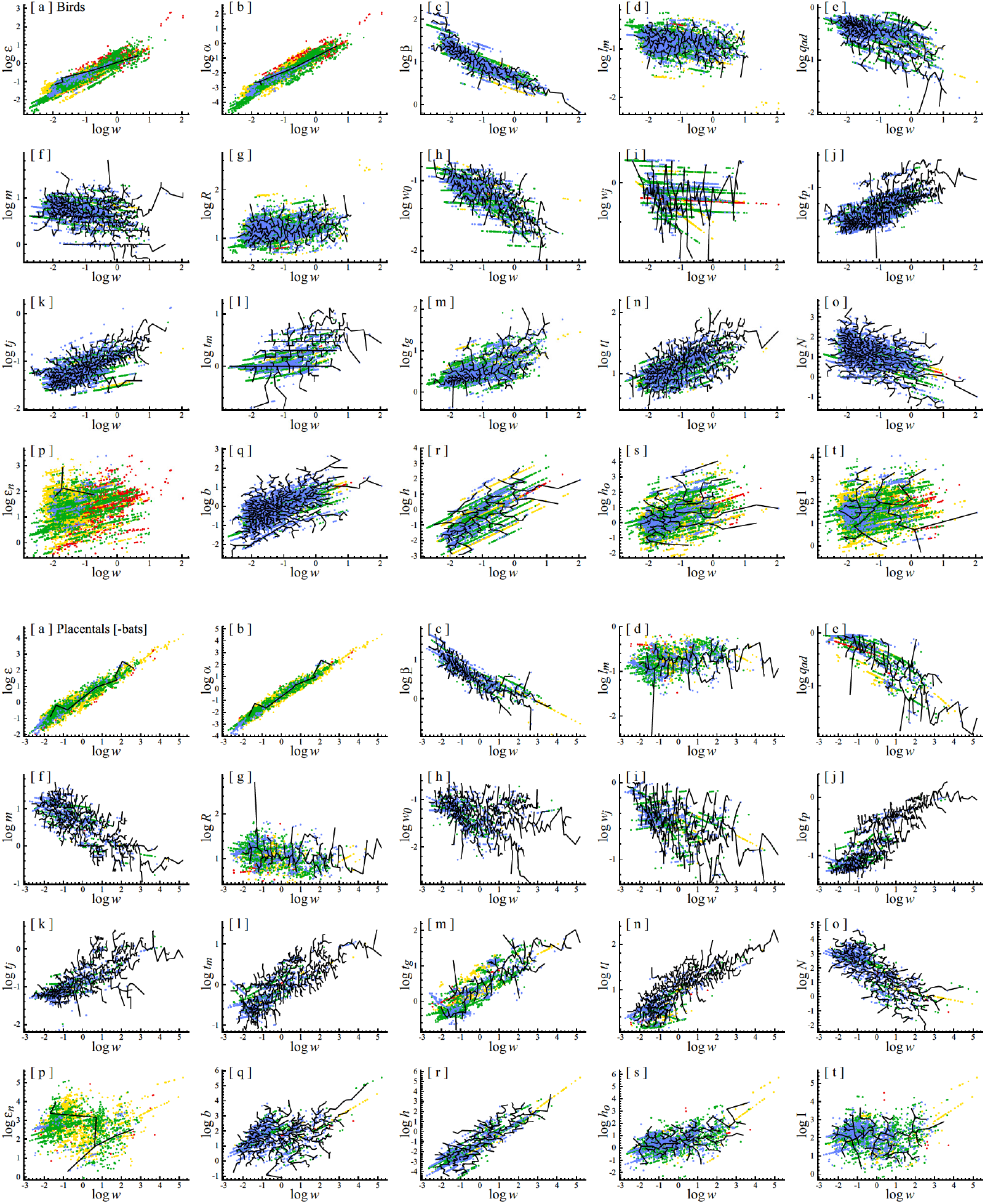
Allometric scaling. Birds in the top four rows, placentals (minus bats) in the bottom four. The non-overlapping black lines connect data, and the coloured dots are estimates at different levels. Estimator levels: data (black), genus (blue), family (green), order (yellow), and class (red); with points of the latter sitting on top of the former.

**Figure 4:**
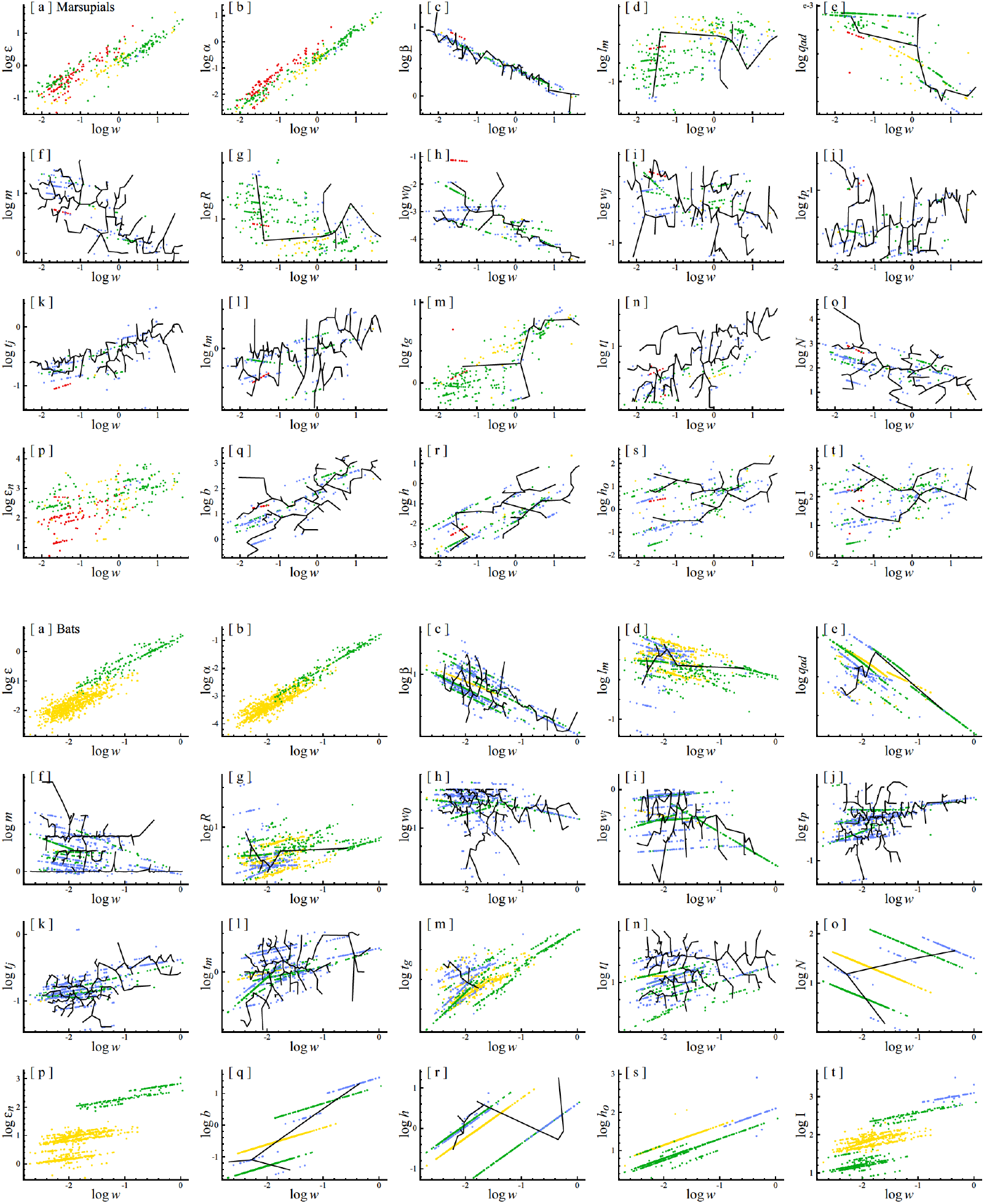
Allometric scaling. Marsupial mammals in the top four rows, bats in the bottom four. The non-overlapping black lines connect data, and the coloured dots are estimates at different levels. Estimator levels: data (black), genus (blue), family (green), order (yellow), and class (red); with points of the latter sitting on top of the former.

### 3.2 Life history variation

Table 4 list the growth and demographic parameters, and Table 5 the energetic and population ecological parameters, for 20 randomly chosen species. The complete set with 11,188 models for birds, and 4,937 models for mammals, are listed in the SI. These estimates illustrate a wide diversity of life histories, with Table 6 summarising the median values of selected traits for birds, placentals (minus bats), marsupials, and bats (BPMB). Table 6 includes also estimates of the allometric intercepts presenting the expected traits of similar sized species of 1kg, and the 95% ranges of the inter-specific variation of the four taxa.

**Table 4:**
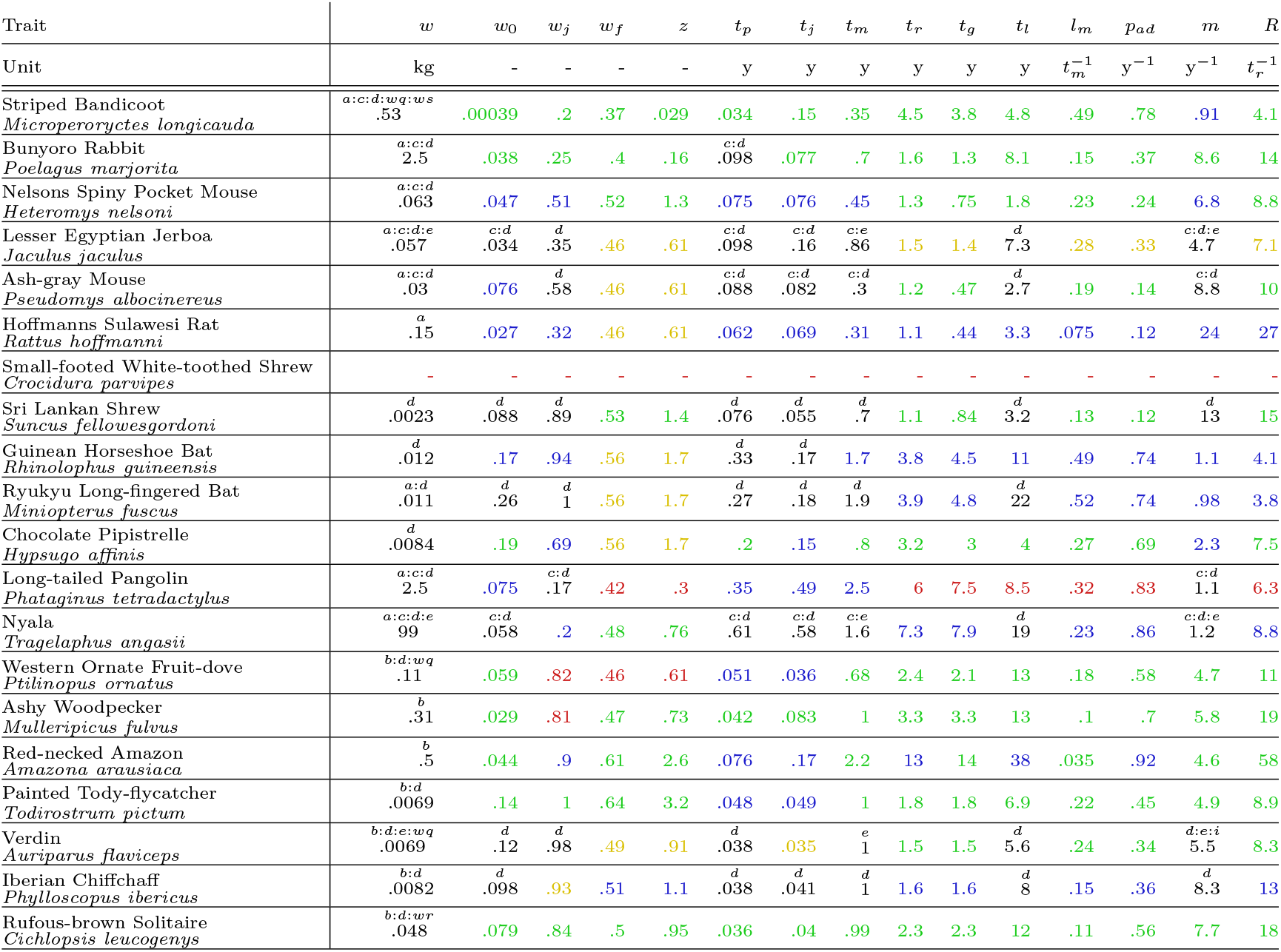
Growth and demography for 20 randomly chosen mammals and birds. Estimator levels: data (black), genus (blue), family (green), order (yellow), and class (red). The superscript letters for data refer to references in the SI on data references.

**Table 5:**
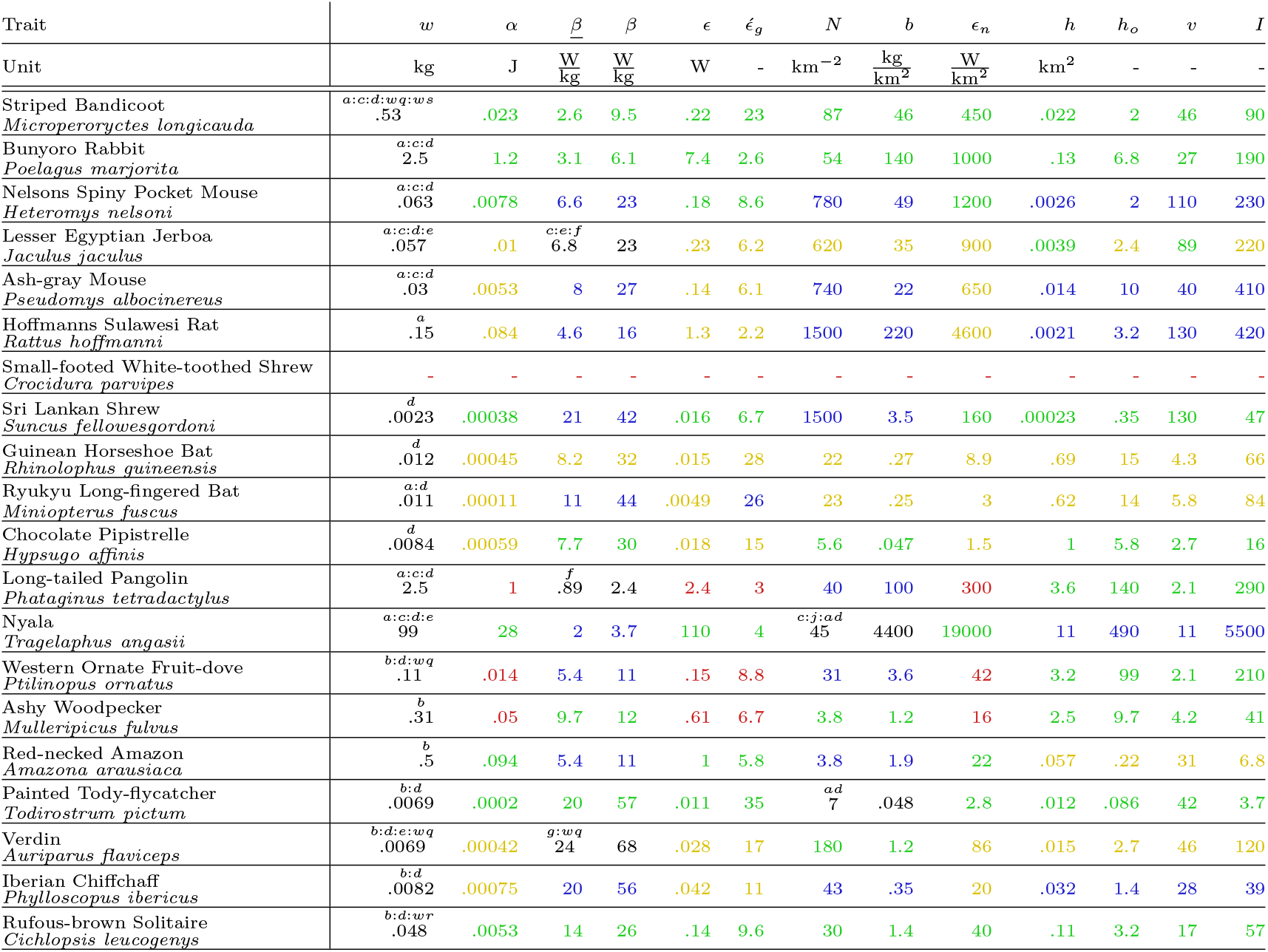
Energetics and population ecology for 20 randomly chosen mammals and birds. Estimator levels: data (black), genus (blue), family (green), order (yellow), and class (red). The superscript letters for data refer to references in the SI on data references.

**Table 6:**
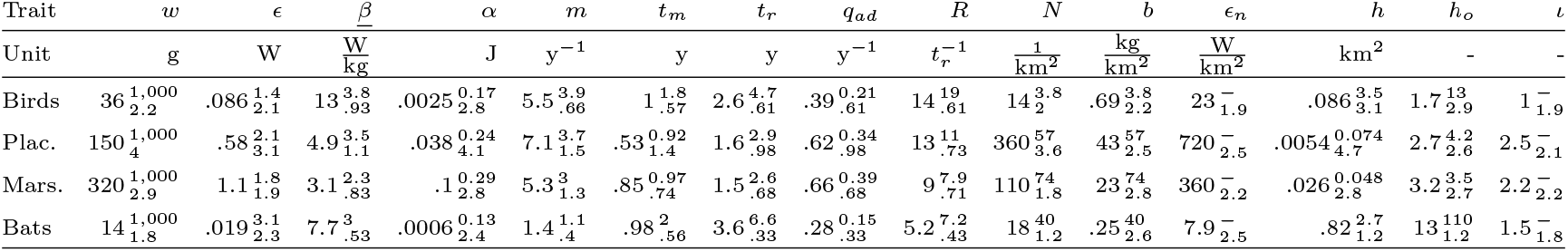
Inter-specific medians of selected traits covering all models for birds, placentals (Plac.), marsupials (Mars.), and bats. Superscript values are allometric intercepts listing expected trait values for body masses of 1kg, and subscripts are the order of magnitudes of the 95% ranges of the inter-specific distributions (the latter calculated for *I* = *e*^*℩*^ for the level of interference).

#### Body mass & net energy

The trait variation across and within the four taxa reflects underlying variation in the conditions of natural selection, with the selected inter-specific range in body mass covering 4.8, 7.9, 4, and 2.7 orders of magnitude for BPMB. The median marsupial is 113% larger than the median placental that is 317% larger than the median bird that is 157% larger than the median bat.

This body mass variation is selected primarily from underlying variation in net energy, where the population dynamic feedback selects net energy from reproduction to body mass—and vice versa—to maintain a constant level of interference competition in the population. This selection predicts the dependence of mass (*w*) on net energy (*ϵ*) as an allometric relation where *w ∝ ϵ*^4*/*3^ (Witting 2017a), with the proportional relation being between body mass and net energy on the per-generation timescale of natural selection.

The *E*^4*/*3^*/w* ratios of the expected 95% range in body mass (calculated from the *ϵ*^4*/*3^ allometry given the 95% range in net energy) over the observed 95% body mass range are 1.22, 1.13, 0.81, and 1.68 for BPMB. This lack of a 100% match is expected because additional body mass variation is selected from allometrically uncorrelated variation in the survival of adults and offspring.

The efficiency by which a species allocates net energy into body mass is reflected in the *w/ϵ* ratio. The medians of this ratio are 0.42, 0.26, 0.29, and 0.74kg/W for BPMB, showing that bats are selected for a more efficient conversion of net energy to mass relative to the other taxa. Marsupials and birds are medium efficient, and placentals least efficient. We will take a closer look at reasons for these differences when we examine the demographic traits below but let us first examine how the different taxa generate net energy.

With the net energy of a species following from the *ϵ* = *αβ* product between resource handling (*α*) and metabolic pace (*β*), the inter-specific variation in net energy and body mass reflects underlying variation in resource handling and mass-specific metabolism. As mass-specific metabolism in birds and mammals approach Kleiber (1932) scaling with allometric exponents around −1*/*4, the 95% range of mass-specific metabolism should cover approximately 25% of the corresponding range in body mass. With the actual *β/w* ratios of the 95% range being 0.42, 0.28, 0.29, and 0.29 for BPMB this holds approximately for most taxa, except that the metabolic estimates for bats have more variation than expected from Kleiber scaling alone.

As the −1*/*4 power decline in mass-specific metabolism with an increase in mass is selected as a secondary mass-rescaling that is imposed by the selection of mass (Witting 2017a), the primary contributor to the selected variation in body mass is the underlying variation in resource handling, reflecting variation in the ways that species handle their resources as well as variation in the density of resources. This downscaling of mass-specific metabolism (*β* ∝ *w*^−1*/*4^) during the selection of mass generates a decline in net energy in physical time (*E* = *αβ* ∝ *w*^−1*/*4^ given constant *α*), but it maintains net energy on the per-generation timescale of natural selection (*ϵt*_*r*_ ∝ *w*^0^ given constant *α*) because it includes a corresponding dilation (*t*_*r*_ ∝ 1*/β* ∝ *w*^1*/*4^) of natural selection time. Thus, when other things are equal, the body mass within a taxon is selected in proportion with resource handling (*w* ∝ *α*).

The dependence of mass on resource handling is reflected in the *α/w* ratio, with the ratios of the 95% ranges of the inter-specific distributions being 1.27, 1.02, 0.97, and 1.33 for BPMB. Except for birds, these ratios are closer to the other-things-being-equal-value of unity than the corresponding *ϵ*^4*/*3^*/w* ratios for net energy given above. The larger *α/w* ratio for birds reflects—at least to some degree—a smaller (−0.33; Witting 2023) than expected (−1*/*4) allometric exponent for mass-specific metabolism. This stronger decline relative to Kleiber scaling implies an additional decline in the pace of handling with increased mass, implying a stronger than proportional dependence of mass on resource handling as the net energy that defines mass is given by the *αβ* product.

The stronger metabolic scaling in birds seems to reflect a diversification that increases as a taxon evolves through time (Witting 2018, 2023), with the primary selection of metabolism accelerating more in smaller species as they are selected over a larger number of generation than larger species. This deviation from a linear metabolic allometry is found also in terrestrial placentals (Kolokotrones et al. 2010), but not in marsupials which have close to perfect Kleiber scaling with an allometric exponent of −0.25 (MacKay 2011). This agrees with the above *α/w* ratio of unity for marsupials, while the close to unity value for placentals is confounded as it is a joint measure for both two-dimensional (2D) and three-dimensional (3D) ecological systems, with the estimated metabolic exponent of placentals being −0.29 for 2D and −0.14 for 3D (Witting 2023). With a metabolic exponent of −0.21, the large *α/w* ratio of 1.33 for bats does not reflect the allometric scaling of metabolism, although it may reflect the larger than expected amounts of metabolic variation in bats.

To examine the different ways the different taxa generate net energy, let us compare the median *β/α* ratios. These are 5200, 130, 31, and 13000/kg s for BPMB, which means that metabolic pace is much more important for the generation of net energy in bats and birds than in placentals and marsupials. If we correct for mass and compare the corresponding *β/α* ratios at the allometric intercept, we find a less extreme but similar ranking (22, 15, 7.9, & 23). The latter reflects a resource handling in bats and birds at 1kg that are only 54% and 71% of the corresponding resource handling in placentals, with mass-specific metabolism in birds and bats at 1kg being 109% and 86% of the corresponding metabolism in placentals. The low importance of metabolism for the generation of net energy in marsupials reflects both a mass-specific metabolism that is 66% of the metabolism in placentals at the 1kg intercept, and a marsupial resource handling at 1kg that is 121% of handling in placentals.

#### Demography

With demographic parameters like *m, t*_*m*_, *t*_*r*_, and *q*_*ad*_ being yearly rates or ages/periods they scale inversely with mass-specific metabolism with allometric exponents for terrestrial birds and mammals around 1*/*4 (Witting 1995, 2023). The 95% range of the inter-specific variation are thus as expected about similar to the corresponding range for mass-specific metabolism, with the ratio of the average range for *m, t*_*m*_, and *t*_*r*_ = 1*/q*_*ad*_ over *β* being 0.66, 1.2, 1.1, and 0.81 in BPMB.

The natural selection of an efficient conversion of net energy into body mass is to a large degree determined by the survival of offspring and adults. With body mass reflecting the available net energy per offspring, the efficiency is proportional to the *t*_*r*_*ϵ/R* ratio of lifetime net energy (*t*_*r*_*ϵ*) over lifetime reproduction (*R*). And with lifetime reproduction being inversely related to the survival of offspring to maturity (*R* = 2*/l*_*m*_ from *λ* = *l*_*m*_*R/*2 = 1), and the reproductive period (*t*_*r*_ = 1*/q*_*ad*_) being inversely related to adult mortality, the efficiency declines if offspring and/or adult mortality increases. A low offspring survival implies that more offspring are produced from the same amount of energy, and a low adult survival implies that less energy is available for the production of the number of offspring produced per lifetime.

These relationships are evident as contrasts in the life history differences between the four taxa. With median offspring survival (*l*_*m*_) in bats being 219%, and median adult survival 162%, of the average median offspring and adult survival in birds, placentals, and marsupials, bats have by far the most efficient conversion of net energy to mass, with a median *w/ϵ* conversion ratio of 229% relative to the average median across the other three taxa. This is reflected in the lowest median lifetime reproduction of 5.2*/t*_*r*_, the longest median reproductive period of 3.6y, and the lowest median annual reproduction of 1.4/y.

Following bats, birds and marsupials have about similar median *w/ϵ* conversion efficiencies of 0.42 and 0.29kg/W, being on average 137% of the median efficiency of placentals. Yet, the ways of obtaining these median conversion efficiencies differ. Where adult bird survival is 179% of adult survival in marsupials at the median, the median survival of marsupial offspring to reproductive maturity is 157% of offspring survival in birds. This implies similar annual reproduction in the two taxa (medians of 5.5/y and 5.3/y), but a low median lifetime reproduction of 9*/t*_*r*_ in marsupials, compared with the largest lifetime reproduction of 14*/t*_*r*_ obtained by birds during their relatively long median reproductive period of 2.6y following low adult survival. Having low median adult survival like marsupials, and low median offspring survival like birds, placentals have the smallest efficiency when it comes to converting net energy into mass.

### Ecology

For the ecological traits there are only few data for bats, and I will thus not include them in the comparison. For the remaining three taxa let us examine the 95% range for abundance (*N*), biomass (*b*), home range (*h*), and home range overlap (*h*_*o*_) relative to the expected range from the variation in body mass given the allometric relations *N* ∝ *w*^−3*/*4^, *b* ∝ *w*^1*/*4^, *h* ∝ *w*^1^, & *h*_*o*_ ∝ *w*^1*/*4^ (I do not include _*n*_ and *℩* as these are predicted invariant of *w*). These ratios show that the average observed range in *N, b, h*, and *h*_*o*_ are 1.06, 1.91, 1.18, and 2.10 times the expected range. These numbers indicate a large overdispersal in *b* and *h*_*o*_ relative to the allometric expectations, which may reflect not only additional ecological variation but also increased estimation uncertainty for ecological traits.

When it comes to between taxa differences there is for most ecological traits a very large difference between birds and mammals (especially placentals). The median abundance, biomass, and resource consumption of placental populations are 26, 62, and 31 times the corresponding medians for birds, and the median home range of placentals is only 6.3% of the median of birds. These differences remain pronounced when corrected for mass, where the median intercepts for abundance and biomass in placentals are 15 times the medians for birds, while the median intercept for placental home range is only 2.1% of the median intercept in birds. These large ecological differences between birds and mammals are in sharp contrasts to the median home range overlap, which is essentially identical between the two taxa (1.7 in birds vs. 2.7 in placentals).

The relatively similar home range overlap in birds and mammals indicates that the spatial packing of home ranges plays a stable central role in the joint natural selection of the life histories and population dynamic ecology as a whole. This is in line with population dynamic feedback selection, where it is the density dependent trade-off between the local resource exploitation of individuals and the interactive competition between individuals that selects a home range overlap that optimises the individual foraging ecology, with the spatial dimensionality of the ecological packing of home ranges constraining the exponents of body mass allometries (Witting 1995, 2017a, 2023).

The level of interference competition that is selected by the spatial packing of home ranges is in fact the main selection attractor of population dynamic feedback selection, known as a competitive interaction fixpoint (Witting 1997). This attractor reflects a process where the population dynamic demography, density regulated abundance, and associated foraging ecology are feedback selected to a balance where they generate a level of interactive competition that is invariant of the selected body mass and pre-defined by the selection attractor of the competitive interaction fixpoint.

Dependent upon the overall constraints on the selection of mass, the level of interference competition at the competitive interaction fixpoint may differ (Witting 2002, 2008, 2017b). The ratio in the *℩*-level of interference for the selection attractor of an exponentially increasing body mass over the attractor of a stable mass, e.g., is 7/3 ≈ 2.3 given a two-dimensional packing of home ranges. This is close to the observed values of 2.5 and 2.2 for the placental/bird and marsupial/bird ratios of the median *℩*-estimates in Table 6.

Apart from the elevated interference competition— and associated increase in population abundance, biomass, and population energy consumption—that is necessary for the selection of an exponential increase in body mass, the feedback selection mechanisms for a stable and increasing body mass are similar selecting no obvious difference in the physiology and demographic traits (while differences are selected in the reproducing unit; Witting 2002). Hence, the median ecological differences, and relative demographic similarities, between birds and mammals (excluding bats) indicate that mammals are predominately selected for an increase in mass while birds are selected for a stable mass.

### 3.3 Population dynamics

The 16,125 equilibrium life history models provided by my study are easily extended into population dynamic simulations. This is exemplified in this section, where eight models are used to explain a diverse range of population dynamic timeseries.

The population dynamics of a species depend on the age-structured demography that defines the timescale of the dynamics, and on the population dynamic regulation that defines not only the bounds on abundance but also the properties of the dynamics, let it be monotonic growth, damped to stable population cycles, fluctuating, or chaotic dynamics.

While regulation is relatively easily estimated from timeseries of abundance estimates, this is not the case for the age-structured demography. It is thus essential that the latter is available from other sources, as the life history models provided in this paper. When these models are extended into age-structured population dynamic models with no regulation, they project a stable population in time.

Given widespread evidence for eco-evolutionary dynamics (e.g., Thompson 1998; Sinervo et al. 2000; Hairston et al. 2005; Coulson et al. 2011; Turcotte et al. 2011a,b; Bell 2017; Hendry 2017; Brunner et al. 2019), I include regulation by density dependent competition and regulation by population dynamic feedback selection, following Witting (2000, 2013). As described in Appendix C, this includes the addition of a density regulation parameter (*γ*) that regulates the growth rate as a function of the density dependent environment, and a selection regulation parameter (*γ*_*℩*_) that accelerates the population dynamic growth below the equilibrium abundance and decelerates population growth above. Following Witting (2021), I use *i*) maximum likelihood statistics to estimate the two regulation parameters (and the equilibrium and initial abundance, as well as an initial measure of competitive quality) and *ii*) the Akaike information criterion (AIC, Akaike 1973) to determine if there is additional evidence for a change in equilibrium abundance or catastrophic mortality in a specific year.

The eight examples of fits to timeseries of abundance estimates are shown in Fig. 5. For these, I estimate that about 20% to 80% of total regulation is caused by selection, as measured by the *γ*_*℩*_*/*(*γ* + *γ*_*℩*_) ratio. This generates a cyclic/over-compensatory dynamics that cannot be explained by density regulation alone. The overcompensation is illustrated by the growth of European Otter (*Lutra lutra*, from Sulkava 2006; LPI 2022) in Finland. The Eurasian Pygmy Owl (*Glaucidium passerinum*, from Knaus and Schmid 2022) in Switzerland illustrates that populations that are both density and selection regulated have the ability to decline for longer periods below the equilibrium abundance, and to increase for longer periods above the equilibrium. Pure density regulated populations will typically only decline above the equilibrium and increase below.

**Figure 5:**
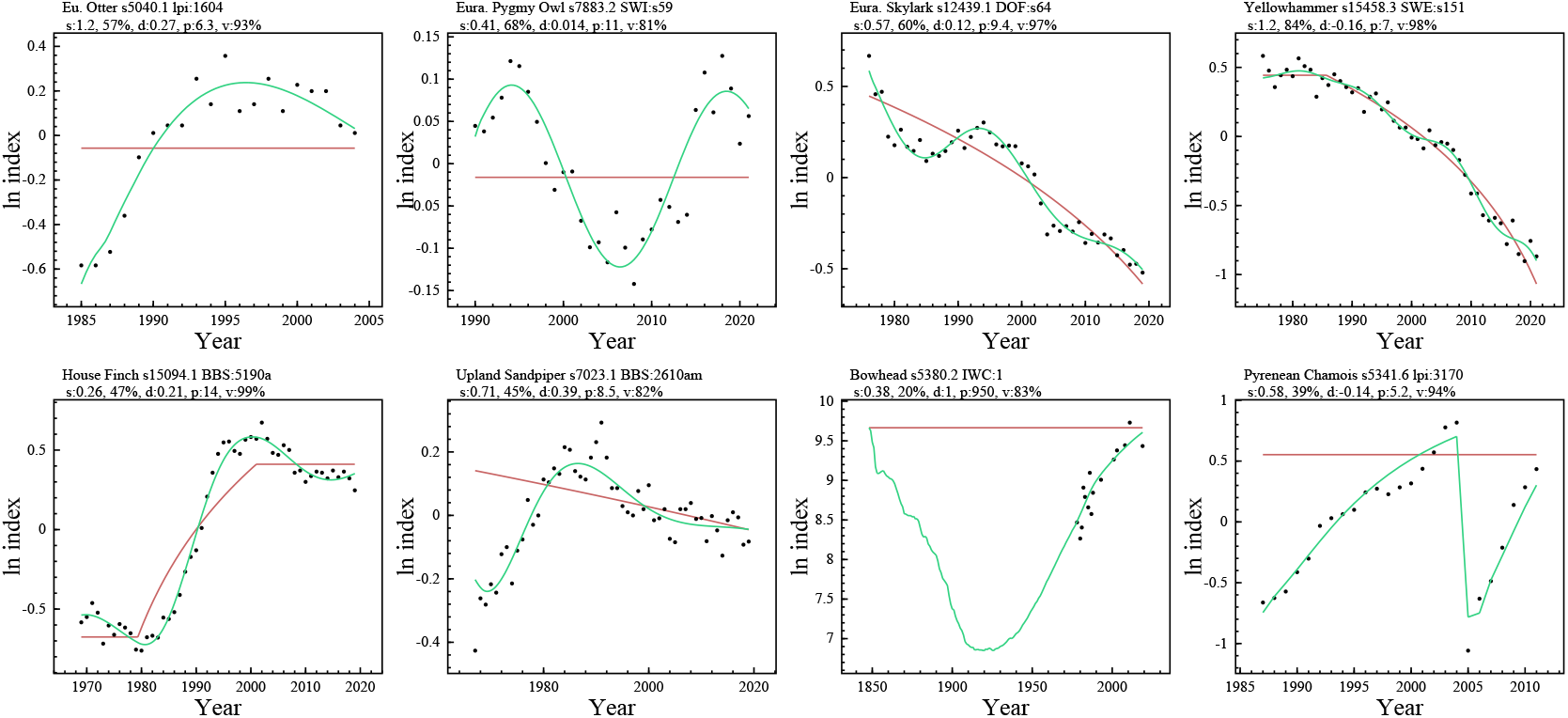
Examples of population dynamic models fitted to timeseries of abundance estimates (for details see the text). Dots are index series of abundance, red lines the estimated equilibria, and green curves the model trajectories. s:*γ*_*℩*_ & *γ*_*℩*_/(*γ*_*℩*_ + *γ*) in %; d:damping ratio; p:period in generations; v:explained variance.

Many birds that live in the farmlands of Europe have declined more or less continuously since systematic monitoring was established in several countries in the 1970th and 1980th (Gregory et al. 2019; PECBMS 2022). This is illustrated by Eurasian Skylarks (*Alauda arvensis*, from DOF 2022) in Denmark and Yellowhammers (*Emberiza citrinella*, from SFT 2022) in Sweden. The best AIC-selected models estimate that Skylark habitats in Denmark and Yellowhammer habitats in Sweden have fragmented and deteriorated by 64% and 80% since the middle of the 1970th.

Where many species suffer, others benefit from habitat changes imposed by humans. Being originally a bird of the western Unites States and Mexico, House Finches (*Haemorhous mexicanus*) established themselves in human-altered habitats throughout the Unites States and southern Canada following a small release on Long Island in the 1940th. Across the mid-west the population was relatively stable from 1970 to 1980, followed by a large increase driven by a three-fold increase in equilibrium (most likely expansion into new habitats), according to the best AIC selected model run on data compiled (Witting 2021) from BBS (2022).

A strong population increase need not necessarily reflect improved conditions. Being a grassland species native to the prairies of Unites States and Canada, the Upland Sandpiper (*Bartramia longicauda*) is another species that is vulnerable to habitats lost to agriculture with an increased use of pesticides. Based on the best AIC selected model run on BBS (2022) data compiled by Witting (2021), from about 1970 to 1990 Upland Sandpipers recovered by an increase of about 50% from a low abundance. Yet, the estimated habitat conditions deteriorated by about 20% from 1970 to 2020, with the species declining by about 20% from 1990 to 2020.

A longstanding threat to many species is direct anthropogenic removals, with commercial whaling in past centuries causing some of the largest perturbations of natural populations worldwide. The population modelling of these perturbations subtracts the historical catches from the simulated population, fitting the model trajectory to a relatively short timeseries of recent abundance estimate (Punt and Butterworth 1999; Wade 2002; Witting 2013). The bowhead whale (*Balaena mysticetus*) in the Bering-Chukchi-Beaufort Seas is one example, where the fitted model estimates that the historical catches of about 21, 000 bowheads from 1850 to 1950 (data from https://iwc.int) caused a collapse to about 1000 individuals in 1920, less than 7% of the pre-whaling abundance. From about 1910 to 1930 there was a turning point where reduced catches and increased growth (from the relaxed density regulation and selection acceleration of the depleted population) turned the population decline into an increase. With bowheads being reproductively mature around 23 years of age, it took approximately 100 years to recover to the pre-whaling abundance of about 15, 000 whales, with todays population continuing to increase.

Population perturbations may also be imposed by natural causes. The Spanish population of Pyrenean Chamois (*Rupicapra pyrenaica*) in Cerdanya-Alt Urgell increased steadily until 2005, where approximately 3,000 died from a border disease virus infection (Marco et al. 2009; data from Lopez-Martin et al. 2013; LPI 2022). This caused the population to collapse by about 80% in a single year. Yet, with Chamois being reproductively mature at about 2 years of age the population was almost recovered by 2011.

## 4 Discussion

Based on 110,105 life history data from 614 literature sources, I used allometric extrapolations at the lowest taxonomic level with data to estimate population dynamic equilibrium life history models for 16,125 species of birds and mammals. With each model having 27 parameters, 435,375 life history and ecological traits were estimated in total. This includes estimates of metabolism, resource handling, net assimilated energy, individual growth, mortality, fecundity, age of reproductive maturity, generation time, life span, home range, population density, home range overlap, biomass, population consumption, and a relative measure of intra-specific interactive competition.

A cross validation of estimates against data confirmed the general absence of estimation bias (Fig. 1), with both allometrically correlated and uncorrelated variation captured by the estimated life history models (Section 3.2). Yet, as the majority of the estimated traits are inter-specific extrapolations, and as the underlying data come from many sources, it is essential to keep in mind the uneven distribution of uncertainties. This is captured to some degree by the removal of outlier data, filter-adjustment of unlikely values, and a colour coding of estimates that reflects the phylogenetic levels of the inter-specific extrapolation, with a separate uncertainty cv calculated for each entry in the trait x estimation-level matrix (Table 3).

Given these limitations, the value of having complete population dynamic life history models for 16,125 species of birds and mammals should not be underestimated. Validated data should always be preferred over model estimates, but life history data are incomplete or missing for most species, and complete life history and population dynamic models are needed in many cases. By aligning my life history estimates with the energy flow in population dynamic feedback selection, it is possible to analyse the life history and population ecological variation in birds and mammals for a better understanding of the natural selection causalities that have created the observed variation. This is illustrated in Section 3.2, where the intra- and inter-taxon variation in birds, placentals, marsupials, and bats are described primarily by the natural selection consequences of variation in resource handling, metabolism, mortality, home range, and abundance. An essential finding suggests that birds are predominately selected for stable body masses, while mammals are predominantly selected for body masses that increase exponentially on evolutionary timescales.

Another example is Witting (2023), where the life history estimates are used to illustrate the natural selection causalities behind the body mass allometries in mammals and birds. This identifies not only the predominant Kleiber scaling, but also allometric deviations imposed by additional variation in mortality and the primary selection of metabolism through time.

The application of the estimated models is not restricted to evolutionary analyse. Another main reason for their development is illustrated in Section 3.3, which shows how the models are easy to extend into complete population dynamic simulations. These examples show how a diverse range of population dynamic timeseries are explained by the addition of population dynamic feedback selection on top of traditional density regulated models. A next step is the estimation of complete population dynamic models across thousands of population dynamic timeseries, allowing for a quantification of the relative importance of regulation by density dependence and natural selection. This is needed for the construction of a freely accessible library of Bird and Mammal Populations (https://mrLife.org/bmp.htm) allowing for online population dynamic analyses of all species.

## 5 Supplementary Information

**si_inflection.pdf:** Data fits of Richards growth curves;

**si_outlier.pdf:** Removal of outlier data;

**si_filter.pdf:** Four estimation filters;

**si_bird demo.pdf:** Bird demography;

**si_bird energy.pdf:** Bird energy/ecology;

**si_mammal demo.pdf:** Mammal demography;

**si_mammal energy.pdf:** Mammal energy/ecology;

**si_ref.pdf:** Data references.

## Acknowledgements

I am grateful to those that collect and published life history and population dynamic data, and to reviewers who improved my study.

## Appendix

### A Demography

I estimate the demography of stable populations with a per-generation growth rate of unity

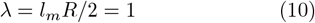

with *l*_*m*_ being the probability that a newborn survives to the age of maturity, and *R* is the expected lifetime reproduction of females that survives to *t*_*m*_, assuming an even sex ratio.

While data on age structured reproduction and survival are becoming increasingly available (e.g., Lemaitre et al. 2020), structured estimates are most often not available, and I estimate lifetime reproduction as a product

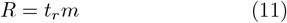

where *t*_*r*_ is the expected reproductive period and *m* the average annual birth rate.

With 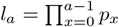 being the survival curve over age (*a*), the expected reproductive period is calculated as a function of adult survival (*p*_*ad*_), where

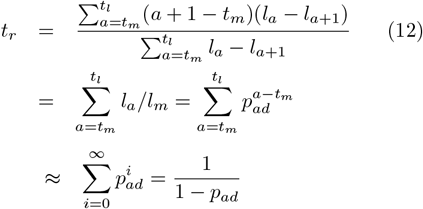

given constant adult survival.

Following Charlesworth (1980), a useful measure of generation time is mean reproductive age

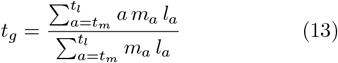

Given constant yearly reproduction (*m*) and constant adult survival (*p*_*ad*_), generation time reduces to a function of the age of maturity and adult survival

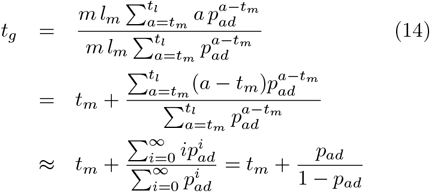

### B Estimation sequence

To estimate life history traits for all species with body mass estimates, I used a sequence of calculation where data and missing parameter estimates are constrained by the underlying demography, growth and energetics. This estimation includes four filters (see filter SI) that use model constraints and data distributions to adjust unexpected estimates for lifespan, adult survival, gross energy, and the growth rated parameters 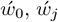 and *t*_*j*_. I converted data on brood size (*m*_*b*_), brood frequency (*m*_*f*_) and/or brood interval (*m*_*i*_) to yearly reproduction (*m* = *m*_*b*_*m*_*f*_, *m*_*f*_ = 1*/m*_*i*_); and data on sexual maturity (*t*_*s*_) to reproductive maturity (*t*_*m*_ = *t*_*s*_ + *t*_*p*_). Data on total metabolism (*βw*) were transformed to metabolic rates per unit mass (*β*), and the ratio (*β/<u>β</u>*) of field (*β*) over basal (*β*) metabolism was calculated for all species with data on both. Mortality (*q*_*ad*_ = 1 − *p*_*ad*_) data were calculated from survival rates (*p*_*ad*_) to have two mirror estimates of all survival data.

Missing values were then estimated by inter-specific extrapolations for *m*, 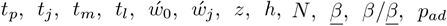 and *q*_*ad*_. Missing values for field metabolism (*β*) were obtained from the *β/<u>β</u>*-ratio and basal metabolism. Missing values for *p*_*ad*_ and *q*_*ad*_ were first estimated independently by allometric correlations on *p*_*ad*_ and *q*_*ad*_, and adjusted afterwards to identical mirror values in order to avoid unrealistic estimates where *p*_*ad*_ *>* 1 ∧ *q*_*ad*_ *<* 0 or *p*_*ad*_ *<* 0 ∧ *q*_*ad*_ *>* 1. With 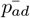 being the average survival rate for the two missing parameter estimates of *p*_*ad*_ and *q*_*ad*_, the adjustment was done by setting *q*_*ad*_ = 1 − *p*_*ad*_ for 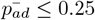, by setting *p*_*ad*_ = 1 − *q*_*ad*_ for 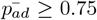, and by setting 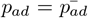 and 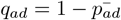 for 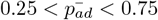.

A survival/mortality filter was constructed from an invariant 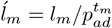 -ratio. As adult survival is larger than offspring and juvenile survival in most species, we expect the *ĺ*_*m*_ ratio to be somewhat smaller than unity in nearly all species. From the *ĺ*_*m*_ distributions across all species, I determined an upper limit on *ĺ*_*m*_ of 0.8 for birds and 0.95 mammals. Then, as *l*_*m*_ = 2*/t*_*r*_*m* = 2(1 − *p*_*ad*_)*/m*, we find that models with *ĺ*_*m*_ ratios above these limits may have an estimate of the reproductive rate (*m*) that is too low, or an estimate of the reproductive age (*t*_*m*_) that is too high, or an estimate of adult survival (*p*_*ad*_) that is too low.

For species that failed to pass the survival filter (i.e., *ĺ*_*m*_ *>* 0.8 for birds; *ĺ*_*m*_ *>* 0.95 for mammals), I would fist examine for the possibility of outlier data for *m* and *t*_*m*_. For this I would, for species with *m* and/or *t*_*m*_ data, estimate the corresponding missing parameter values for *m* and *t*_*m*_. If one of the latter two estimates made the species pass the survival filter, I would classify the data value as an outlier. If no outlier was identified I would increase adult survival to obtain a *ĺ*_*m*_ ratio that resembled the missing parameter estimate of *ĺ*_*m*_ for that species. I would then remove the *p*_*ad*_ and *q*_*ad*_ outliers from the main data and rerun the complete estimation sequence until no extra outliers were identified.

Estimates were then calculated as *t*_*r*_ = 1*/*(1 − *p*_*ad*_) by eqn 12, *t*_*g*_ = *t*_*m*_ + *p*_*ad*_*/*(1 − *p*_*ad*_) by eqn 13, 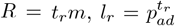, and *l*_*m*_ = 2*/t*_*r*_*m* from zero growth *λ* = *l*_*m*_*t*_*r*_*m/*2 = 1 at equilibrium.

A lifespan filter adjusted unlikely small *t*_*l*_ values, and all growth related parameters 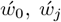, and *t*_*j*_ were filtered for unexpected values. 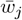 was then estimated by eqn 6, net energy as 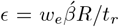, gross energy as _*g*_ = *wβ* + */*2 including the gross energy filter, resource handling as *α* = */β*, home range overlap as *h*_*o*_ = *hN*, encounter rate as *v* = *<u>β</u>w*^1*/*2*d*^*/h*^1*/d*^, and the level of interference as *I* = *h*_*o*_*v* followed by a rescaling to set *℩* = ln *I* equal to unity at the median across all birds.

### C Population dynamics

The age-structured population dynamic model is parameterised from the age of reproductive maturity in years (*t*_*m*_), annual reproduction (*m*), and annual survival (*p*_*ad*_) of adults. These parameters are converted to an appropriate timescale where the timestep of the iteration is Δ*t* ≤ min(1, *t*_*m*_), with parameters scaled as *a*_*m*_ = *t*_*m*_*/*Δ*t, m*^∗^ = *m*Δ*t*, and *p* = *p*_*ad*_^1*/*Δ*t*^.

The survival parameter *p* is used as the survival rate for all age-classes except age-class zero. To calculate age-class zero survival, I converted *p* into the reproductive period *t*_*r*_ = 1*/*(1 − *p*), to calculate lifetime reproduction *R* = *t*_*r*_*m*^∗^, and *l*_*m*_ = 2*/R* from the equilibrium constraint *l*_*m*_*R/*2 = 1. Age class zero survival is then calculated as 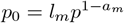.

With *x* " *a*_*m*_ being the maximum lumped age-class, the number *n*_*a,t*_ of individuals of age 0 *< a < x* at timestep *t* is

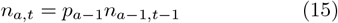

and the number in age-class *x*

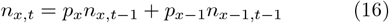

with *p*_*a*_ = *p*_0_ for *a* = 0 and *p*_*a*_ = *p* for *a* ≥ 1. Let the number of individuals in each age-class relate to time just after each timestep transition, with offspring at *t* being produced by the *t* − 1 individuals that survive to the *t* − 1 → *t* transition, with the density dependent ecology being approximated by the average 1+ abundance of the two timesteps:

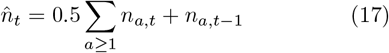

The number of offspring in age-class zero is then

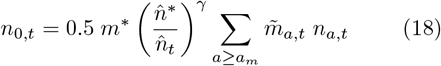

where *γ* is the strength of density regulation, and 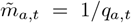, with *q*_*a,t*_ being the average competitive quality of cohort *a* at *t*. At the population dynamic equilibrium 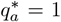 for all ages. More generally *q*_*a,t*_ = *q*_*a*−1,*t*−1_ and

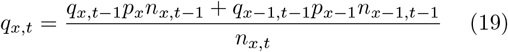

assuming that there is no change in the quality of a cohort over time. The quality of offspring

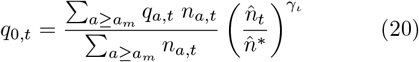

is the average quality of the mature component multiplied by the density dependent selection, with *γ*_*℩*_ being the selection response.

For the initial conditions of an iteration I use the same quality across all individuals, and an initial abundance with a stable age-structure

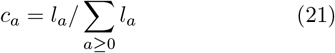

where *l*_0_ = 1, *l*_*a*_ = *p*_0_ *p*^*a*−1^ for 1 ≤ *a < x*, and *l*_*x*_ = *p*_0_ *p*^*x*−1^*/*(1 − *p*).

